# Characterization of glutamatergic VTA neural population responses to aversive and rewarding conditioning in freely-moving mice

**DOI:** 10.1101/464040

**Authors:** Quentin Montardy, Zheng Zhou, Zhuogui Lei, Xuemei Liu, Pengyu Zeng, Chen Chen, Yuanming Liu, Paula Sanz-Leon, Kang Huang, Liping Wang

**Affiliations:** Shenzhen Key Lab of Neuropsychiatric Modulation and Collaborative Innovation Center for Brain Science, CAS Center for Excellence in Brain Science and Intelligence Technology, the Brain Cognition and Brain Disease Institute (BCBDI) for Collaboration Research of SIAT at CAS and the McGovern Institute at MIT, Shenzhen Institutes of Advanced Technology, Chinese Academy of Sciences, Shenzhen 518055, China; University of Chinese Academy of Sciences, Beijing 100049, China; School of Physics, The University of Sydney, Sydney 2006, Australia; Centre for Integrative Brain Function, The University of Sydney, Sydney 2006, Australia

**Keywords:** Ventral Tegmental Area, Aversion, Reward, Conditioning, Behavior

## Abstract

The Ventral Tegmental Area (VTA) is a midbrain structure known to integrate aversive and rewarding stimuli, a function involving VTA Dopaminergic and GABAergic neurons. VTA also contains a less known population: glutamatergic (VGluT2) neurons. Direct activation of VGluT2 soma evokes rewarding behaviors, while stimulation of their axonal projections to the Nucleus Accumbens (NAc) and the Lateral Habenula (LHb) evokes aversive behaviors. Here, a systematic investigation of the VTAV^GluT2+^ population response to aversive or rewarding conditioning facilitated our understanding these conflicting properties. We recorded calcium signals from VTA glutamatergic population neurons using fiber photometry in VGluT2-cre mice to investigate how the VTA glutamatergic neuronal population was recruited by aversive and rewarding stimulation, both during unconditioned and conditioned protocols. Our results revealed that, as a population, VTA^VGluT2+^ neurons responded similarly to unconditioned-aversive and unconditioned-rewarding stimulation. During aversive and rewarding conditioning, the CS-evoked responses gradually increased across trials whilst the US-evoked response remained stable. Retrieval 24 h after conditioning, during which mice received only CS presentation, resulted in VTAV^GluT2+^ neurons strongly responding to CS presentation and to the expected-US but only for aversive conditioning. The inputs and outputs of VTA^VGluT2+^ neurons were then investigated using Cholera Toxin B (CTB) and rabies virus, and we propose based on all results that VTAV^GluT2+^ neurons specialized function may be partially due to their connectivity.

## Introduction

### The Ventral Tegmental Area

The ventral tegmental area (VTA) is a midbrain structure that has been linked with a variety of behavioral functions including aversion and reward (Sanchez-Catalan et al., 2014; Bromberg-Martin, Matsumoto, and Hikosaka, 2010; Lammel, Lim, and Malenka, 2014; Barker et al., 2016), prediction error (Watabe-Uchida, Eshel, and Uchida, 2017; Bromberg-Martin, Matsumoto, and Hikosaka, 2010), and motivation (Arsenault et al., 2014; Bromberg-Martin, Matsumoto, and Hikosaka, 2010; van Zessen et al., 2012). The heterogeneous composition of VTA includes a large proportion of dopaminergic (DA) neurons (60%), and smaller proportions of GABAergic neurons (GABA) (35%) and glutamatergic neurons (2-5%) (Yamaguchi, Sheen, and Morales, 2007; Yamaguchi et al., 2015; Nair-Roberts et al., 2008). Whilst the two first neuronal components (DA and GABA) of VTA have been well studied and characterized (Morales and Margolis, 2017; Sanchez-Catalan et al., 2014), the glutamatergic population has received less attention and its functional characterization needs to be elucidated to obtain a better understanding of VTA function.

### VTA dopaminergic and GABAergic neuronal function during aversion and reward

In-vivo electrophysiology experiments have shown that VTA dopaminergic (DA) neurons increase their firing rate following rewarding stimulation (Lammel, Lim, and Malenka, 2014; Schultz, Dayan, and Montague, 1997; Bromberg-Martin, Matsumoto, and Hikosaka, 2010). However, if the reward is paired with a conditioned stimulus (CS), such as a sound cue, the activity of DA neurons gradually shifts from responding to the reward presentation, to responding to the CS presentation (Bromberg-Martin and Hikosaka, 2009; Bromberg-Martin, Matsumoto, and Hikosaka, 2010). At the same time, VTA GABAergic neurons can signal expected outcome (Cohen et al., 2012) by inhibiting neighboring DA neurons (Cruz et al., 2004; Tan et al., 2010; Tan et al., 2012; Eshel et al., 2015), which corresponds with the decrease in firing rate observed in DA neurons following aversive stimulation (Ungless, Magill, and Bolam, 2004). This mechanism of reward prediction via VTA GABAergic neuronal inhibition of local DA neurons is further supported by recent optogenetic experiments (Tan et al., 2012), in which direct optogenetic activation of VTA-GABA neurons leads to local VTA-DA neuronal inhibition and also to place aversion (Tan et al., 2012).

### Physiology, anatomy and function of VTA glutamatergic neurons

While there is a general consensus regarding the role of VTA-DA and VTA-GABA neurons in control of aversive and rewarding behavior, the role of VTA’s glutamatergic neurons is not yet understood (Morales and Root, 2014; Morales and Margolis, 2017). Glutamatergic neurons, defined by their expression of vesicular glutamate transporter 2 (VGluT2) (Yamaguchi, Sheen, and Morales, 2007; Kawano et al., 2006), can form local connections in VTA (Dobi et al., 2010) and send long-range projections to structures such as the Nucleus Accumbens (NAc) or the Lateral Habenula (LHb) (Taylor et al., 2014; Qi et al., 2016; Root et al., 2014; Root et al., 2014). In addition, VGluT2 neurons are a heterogeneous population in terms of molecular and physiological characteristics: one population of VGluT2 neurons releases only the excitatory neurotransmitter glutamate, whilst another fraction can release glutamate and can also co-release dopamine or GABA (Yamaguchi et al., 2011; Morales and Margolis, 2017; Root et al., 2014; Kawano et al., 2006).

Optogenetic stimulation of VTAV^GluT2+^ neuronal somata promotes rewarding behaviors, such as place preference and appetitive instrumental conditioning (Wang et al., 2015); however, at the microcircuit level the function of their interaction with neighboring DA and GABA neurons remains unknown (Dobi et al., 2010). Conversely, VTA glutamatergic transmission has also been associated with aversive behaviors (Han et al., 2017). Indeed, optogenetic activation of axonal projections from VTA^VGluT2+^ neurons mainly elicit aversive responses, such as escape or avoidance. More specifically, optogenetic activation of VTA^VGluT2+^ terminals in the nucleus accumbens (NAc) or to the lateral habenular nucleus (LHb) both promote aversive behaviors, including aversive conditioning (Root et al., 2014; Wang et al., 2015). Recent electrophysiological recordings of glutamatergic neurons confirmed that individual VTA^VGluT2+^ neurons can be activated by aversive stimulation, and either excited or inhibited by rewarding stimulation (Root, Estrin, and Morales, 2018), giving an initial insight into this seemingly paradoxical function. However, population response recordings, such as calcium imaging, can potentially help us to understand VTA^VGluT2+^ function in terms of conditioned aversion and reward. In particular, we use fiber photometry with the genetically-encoded Ca^2+^ indicators GCaMP6s (Chen et al., 2013), which is a minimally invasive method that allows in-vivo measurements in freely-moving animals of synchronous neuronal population activity from subcortical structures (Resendez and Stuber, 2014; Guo et al., 2015), which has proved useful during conditioning (Daqing et al., 2017).

Here, we investigated the VTA^VGluT2+^ neuronal population response to aversive and rewarding events, both during unconditioned and conditioned stimulation. Conditioning consisted in pairing a tone (CS+) with an aversive (footshock) or rewarding (sucrose) unconditioned stimulus (US). In addition, a retrieval test was performed 24 h after conditioning to see whether VTA^VGluT2+^ neurons maintain a robust memory of aversive or rewarding conditioning. Finally, to better understand VTA^VGluT2+^ population responses to aversion and reward, we investigated their connectivity pattern using cell specific monosynaptic retrograde rabies virus tracing, allowing the mapping VTA^VGluT2+^ inputs.

## Materials and Methods

### Animals

All procedures were approved by Animal Care and Use Committees in the Shenzhen Institute of Advanced Technology (SIAT) or Wuhan Institute of Physics and Mathematics (WIPM), Chinese Academy of Sciences (CAS). Adult (6-8 weeks old) male VGluT2-ires-cre (Jax No.016963, Jackson Laboratory) transgenic mice were used in this study. All mice were maintained on a 12/12-h light/dark cycle at 25°C. Food and water were available ad libitum.

### Viral preparation

For fiber photometry experiments, AAV2/9-EF1a-DIO-Gcamp6s virus was used. For tracing experiments, we used AAV2/9-EF1a-DIO-BFP. Virus titers were approximately 2-3×1012 vg/ml. In the rabies tracing experiments, AAV2/9-CAG-DIO-histone-TVA-GFP (4.2×1012 vg/ml), AAV2/9-CAG-DIO-RV-G (4×1012 vg/ml) and EnvA-RV-DsRed (1×109 pfu/ml) viruses (BrainVTA Co., Ltd., Wuhan) were used.

### Viral injections

VGluT2-ires-cre mice were anesthetized with pentobarbital (i.p., 80 mg/kg) and fixed on stereotaxic apparatus (RWD, Shenzhen, China). During the surgery, mice were kept anesthetized with isoflurane (1%) and placed on a heating pad to keep the body temperature at 35°C. A 10 μl microsyringe with a 33-Ga needle (Neuros; Hamilton, Reno, USA) was connected to a microliter syringe pump (UMP3/Micro4; WPI, USA) and used for virus injection into VTA (coordinates: AP, –3.15mm; ML, –0.3 mm; DV,–4.4mm).

### Retrograde tracing

For CTB tracing, VGluT2-ires-cre mice received CTB Alexa Fluor conjugates (CTB 594 or CTB 488, Invitrogen Inc., Grand Island, NY, USA) via injection into NAc and LHb (50 nl per injection) and AAV2/9-EF1a-DIO-BFP into VTA (200 nl per injection). For trans-synaptic rabies tracing, a total volume of 70 nl mixed viruses AAV-EF1a-DIO-RV-G and AAV-EF1a-FLEX-GFP-TVA (volume ratio: 1:1) were injected into VTA of VGluT2-ires-cre mice (coordinates: AP, –3.15mm; ML, –0.3 mm; DV,–4.4mm). After 3 weeks, 100 nl of EnvA-RV-DsRed (EnvA-pseudotyped rabies virus) was injected at the same coordinates. Mice were sacrificed one week after this second injection. All rabies tracing experimental procedures were completed in Biosafety level 2 (BSL2) Laboratory.

### Implantation of optical fibers

A 200 μm optical fiber (NA: 0.37; NEWDOON, China) was chronically implanted in the VTA of VGluT2-ires-cre mice 2-3 weeks following virus expression for fiber photometry experiments. The optical fiber was unilaterally implanted in VTA (AP:–3.15 mm, ML: –1.10 mm and DV:–4.2 mm) with a 15° angle in the medial direction of the transverse plane. After surgery all mice were allowed to recover for at least 2 weeks.

### Histology, immunohistochemistry, and microscopy

Mice were sacrificed by overdosing with pentobarbital (1% m/v, 150 mg/kg, i.p.) and transcardially perfused with 1M cold saline followed by ice-cold 4% paraformaldehyde (PFA; Sigma) in 1M PBS. Brains were removed and submerged in 4% PFA at 4°C overnight to post-fix, and then transferred to 30% sucrose to equilibrate. The coronal brains slices (40 μm) were sectioned with a cryostat (CM1950; Leica, Germany). Freely floating sections were washed with PBS and blocked for 1 h at room temperature in blocking solution containing 0.3% Triton X-100 and 10% normal goat serum (NGS). Then the sections were incubated overnight with rabbit monoclonal anti-dsRed (1:500, #632496; Clontech; Japan); GFP(1:500, #ab290, abcam, USA); DAPI (1:50,000, #62248; Thermo Fisher Scientific, USA) diluted in PBS with 3% NGS and 0.1% TritonX-100. The sections were incubated for 1 h at room temperature with Alexa Fluor 488 or 594 goat anti-rabbit secondary antibody (1:200; Jackson Laboratory, USA). Finally, the sections were mounted and photographed using the Zeiss LSM 880 confocal microscope (Zeiss; Germany). The images were acquired using identical gain and offset settings, and analyzed with ImageJ, Image Pro Plus, and Adobe Photoshop software. ROIs were traced with reference to the “The mouse brain in stereotaxic coordinates” by George Paxinos and Keith B. J. Franklin. CTB and Rabies Virus immunoreactivity was quantified using Image Pro Plus and was verified by comparing with manual counts performed by a trained double-blind observer.

### Unconditioned aversive and Conditioned aversive stimulation

VGluT2-ires-cre mice (N=8) with optical fibers implanted were placed in an unescapable acrylic box (L 25 × W 25 × H 70 cm) with a metal grid floor that delivered footshock currents (0.6 mA footshock, 0.5 s). Each mouse went through an unconditioned and then a conditioned protocol as below. During unconditioned aversive stimulation, mice were freely moving and footshocks were directly delivered with inter-trial interval durations varying within session randomly set in a range between 60-120 s. The session was approximately 10 min long and each mouse received 10 footshoocks. The conditioned sessions consisted of 5 trials where an auditory conditioned stimulus (CS; 3 kHz, sine wave, 90 dB, 5 s) was paired with an unconditioned stimulus (US; 0.5 s, 0.6 mA footshock; random intertrial intervals 60-120 s) that began immediately after tone ended. Mice were presented with 5 CS cues alone, without footshock stimulation, 24 h after conditioning.

### Unconditioned Reward and Conditioned reward

Following viral injections and optical fiber implantations, VGluT2-ires-cre mice (N=8) underwent a third surgery to implant a steel headplate for head-fixing purposes (Chen et al., 2013). The mice were habituated (~30 min/day) to the head-fix system over two-three days. During the experiment, each mouse was head-fixed and a tube delivering liquid reward was directly aimed at their mouth, through which single drops of sucrose (5% w/v) could be delivered as reward. Each mouse went through an unconditioned and then a conditioned protocol as described below. Unconditioned reward sessions were conducted during which 30 reward trials were presented with inter-trial interval durations varying within session, randomly set in a range between 25-40 s. Conditioned reward sessions consisted of one session of 30 trials in which an auditory conditioned stimulus (CS; 10 kHz, sine wave, 80 db, 5 s) was paired with one sucrose delivery, which was delivered immediately after the tone ended. Mice were presented with 30 CS cues alone, without sucrose reward, 24 h after conditioning.

### Fiber photometry

Ca^2+^ signals were recorded using a fiber photometry system (Thinker Tech, Nanjing). Two weeks post AAV2/9-DIO-Gcamp6s virus injection, an optical fiber (NA: 0.37; NEWDOON, China) was implanted into VTA as described above.

The fiber photometry system included a 502–730 nm transmission band (Edmund, Inc.), a 480 nm excitation light from LEDs (CREE XPE), reflected off a dichroic mirror with a 435–488 nm reflection band and coupled into a 200 μm 0.37 NA optical fiber (Thorlabs, Inc.) by an objective lens. At the fiber tip, the laser intensity was about 20 μW. The collection of Gcamp6s fluorescence used the same objective, transmitted by the dichroic mirror filtered through a green fluorescence protein (GFP) bandpass emission filter (Thorlabs, Inc. Filter 525/39), and detected by a CMOS camera sensor (Thorlabs, Inc. DCC3240M). The calcium signals were recorded by CMOS camera at 50 Hz. A LabVIEW (National Instruments, US) program was developed to control the CMOS camera. Behavioral event signals were recorded by a DAQ card (NI, usb-6001) at 1000 Hz using the same LabVIEW program.

### Photometry data analysis

Calcium Imaging signals were first extracted using Blackrock NPKM (Neural Processing MATLAB Kit), using provider instructions (Thinker Tech, Nanjing). Custom MATLAB (The MathWorks Inc. ©) scripts were developed for further analysis using R2012a. Signals were analyzed as dF/F = (F − Fb)/Fb, where Fb was defined as the baseline fluorescence before stimulation. Data were then smoothed using a 10 ms sliding windows. Time courses were calculated by aligning the time of stimulation across all individual trials and then calculating the mean change in calcium at each time window. To compare calcium activity between conditions, mean calcium activity was calculated for 0.5 s time windows centered around the time of the activity peak (2 s before stimulation vs. CS vs. US). A multivariate permutation (1000 permutations, α level of 0.05) test was used to test the statistical significance of the difference between conditions over the time course, and a threshold indicating a statistically significant difference from the baseline was applied (p<0.005). Area Under Curve index is the sum of transient Ca^2+^ activity (Wang, 2017) over a period of 0.5 s centered around the peak of activity.

## Results

### 1. VTA^VGluT2+^ population increases activity to unconditioned aversive stimulation and this response remains constant over successive trials

We first investigated whether the VTA^VGluT2+^ neuronal population responds to unconditioned aversive stimulation. To do that we first infected VGluT2-cre animals with adeno-associated virus (AAV) expressing GCaMP indicator by injecting the AAV9-EF1a-DIO-GCaMP6s virus in VTA (**Fig. 1.A**). Three weeks later an optical fiber was implanted above VTA, allowing in-vivo recording of VTA^VGluT2+^ calcium signals during freely-moving behavior (**Fig. 1.A-B**). At the end of experiments, GCaMP6s virus expression in VTA (**Fig. 1.C**) and fiber positioning were systematically checked in every mouse. (**Fig. 1.D**). During the unconditioned aversive experiment, mice received footshocks (0.5 s at 0.6 mA) whilst the activity of VTA^VGluT2+^ neurons were recorded. Immediately after the beginning of the footshock, the calcium signal of VTA^VGluT2+^ neurons strongly increased for each individual mouse (**Fig. 1.E, top**), which was a stereotypical effect well aligned with the onset of stimulation (**Fig. 1.E, bottom**). All mice expressed a similar increase of activity (4.26% DF/ F, n=8) directly after aversive stimulation, which was significantly different from baseline expression for 1.54 s before returning to baseline level (**Fig. 1.F**, red part of the curve indicates p<0.05 using the multivariate permutation test). The mean signal values for all mice for a period of 0.5 s around the peak response amplitude (T=0.68 s) revealed that activity was significantly higher than baseline (BL=0.002% DF/F vs. Footshock=3.96% DF/ F, p<0.0001; **Fig. 1.G**). To observe the effect of repeated unconditioned stimulation on VTA^VGluT2+^ neurons, we analyzed response trends on a trial-by-trial basis and across animals (**Fig. 1.H**). The peak responses in successive trials remained at a similar level (**Fig. 1.H**), which was confirmed by computation of the Area Under Curve index (AUC) (**Fig. 1.I**).

**Figure 1.**
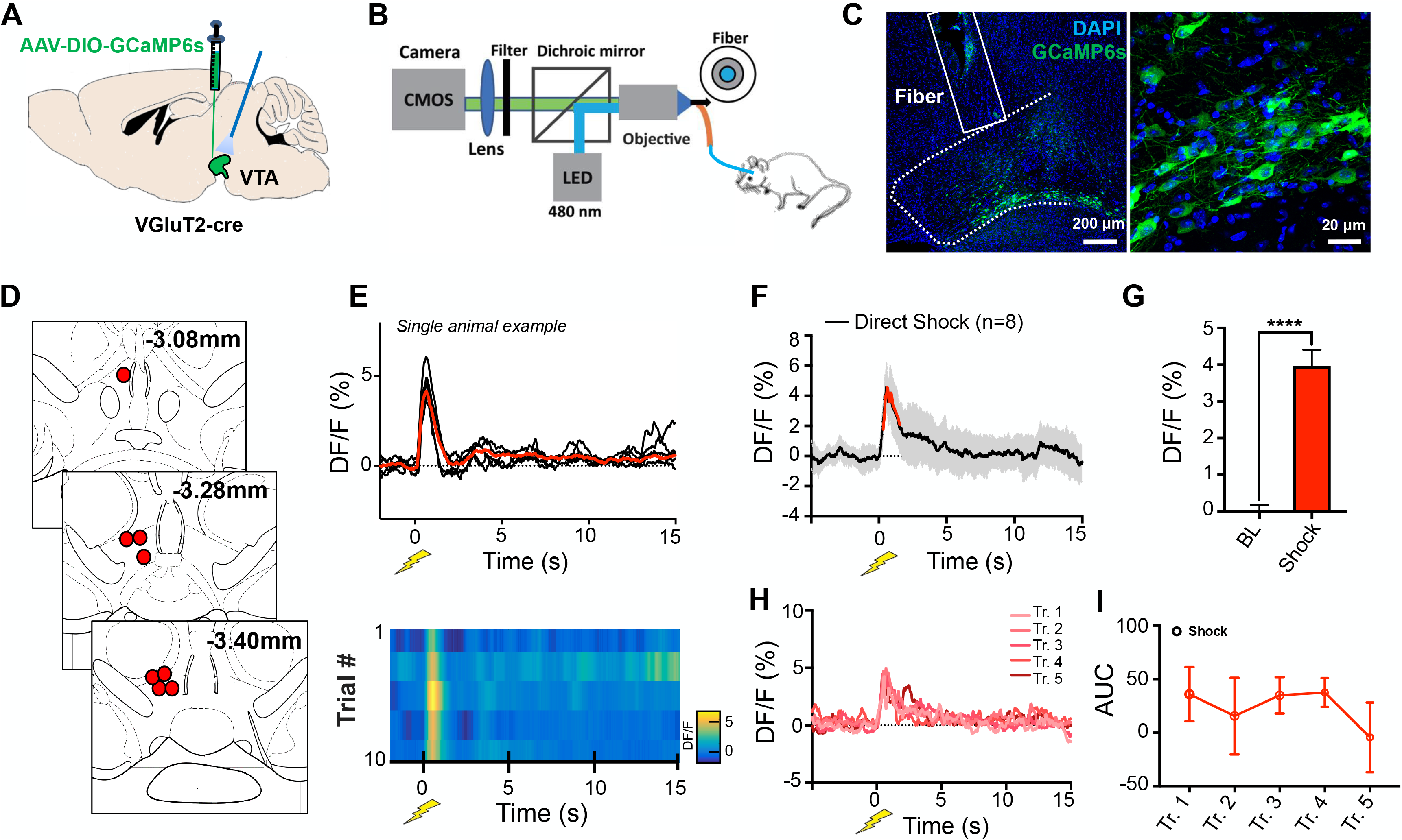
VTA response to unpredicted aversive stimulation. **A.** Schematic representation of AAV9-DIO-GCaMP6s injection in VTA. **B.** Schematic representation of Fiber Photometry setup. **C.** Representative image of GCaMP6s virus expression in VTA of VGluT2-cre mice (Green, GCaMP6s; scale bar, 200 μm and 20 μm respectively). **D.** Location of optical fiber position for every mouse. **E.** Example showing individual trials Ca2+ signal in VTA VGluT2 for one mouse (top) plotted as an individual time courses in black, and averaged in red; (bottom) plotted as heatmap. Footshock stimulation at T=0 ms; Lightning symbol represents footshock time (T=0). **F.** Mean time course of VTA VGluT2+ calcium signal (n=8 mice), increasing during unconditioned stimulation. Data points in red represent a section of the time course where differences compared to base line (BL) are statistically significant, using a Multivariate Permutation Test.**G.** Mean Ca2+ signal during the baseline vs. Shock (BL=0.002% DF/F vs. Shock=3.96% DF/F, P<0.0001). **H.** Mean trial-by-trial responses (trial 1 to trial 5) **I.** Trial-by-trial Area Under Curve (AUC) centered on shock stimulation, computed for all animals.

In summary, this experiment demonstrated that VTA^VGluT2+^ neurons were strongly activated by unconditioned aversive stimulation and the amplitude of the peak response remained constant across trials.

### 2. The VTA^VGluT2+^ neuronal population responds to aversive conditioning

To characterize the responses of VTA^VGluT2+^ to conditioned aversive stimulation (**Fig. 2.A**) we applied the following protocol: (i) Habituation Day, during which animals received tone stimulation only; (ii) Conditioning Day, during which a tone (CS) was paired with an unconditioned stimulation (US); (iii) Retrieval Day, during which only the CS was presented.

**Figure 2.**
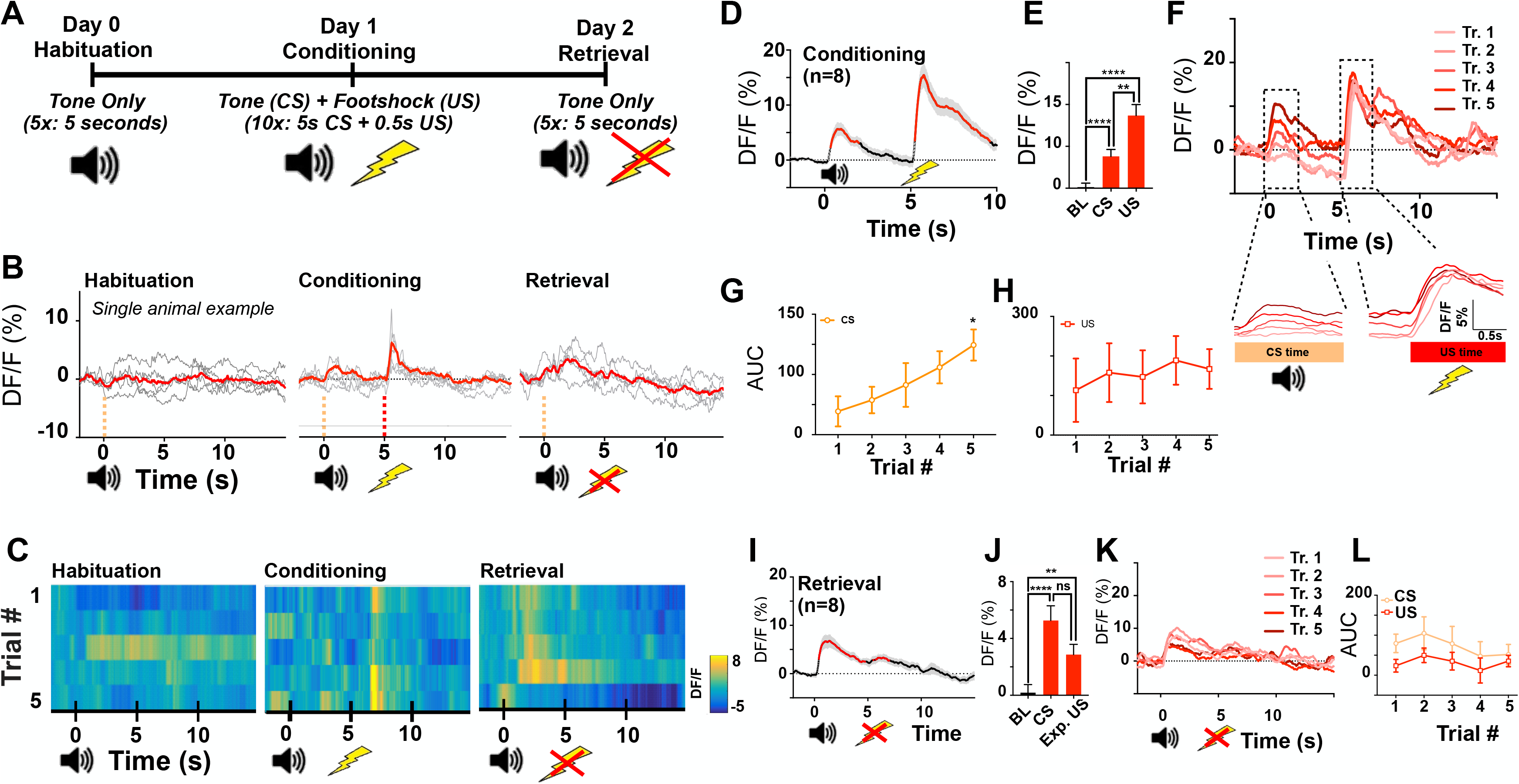
VTA response to aversive conditioning. **A.** Schematic representation of CS-US conditioning, where CS is a 5 s tone and US a 0.5 s footshock starting immediately after CS; Symbols represent tone CS and US footshock time (respectively T=0s and T=5s). **B.** Single animal example signal plotted as individual time courses in black, and averaged in red, for Day 0 Habituation (tone only), Day 1 Conditioning (CS-tone + US Shock), and Day 2 Retrieval (CS only). **C.** Same as panel B, plotted as heatmap. **D.** Day 1 Mean Ca2+ signal over the conditioning time course (n=8 mice), showing an increase during CS and US stimulation. **E.** Mean Ca2+ signal comparing Baseline (BL), CS and US (BL-evoked=0.12%DF/F; CS-evoked=3.8%DF/F, p<0.0001; US-evoked=8.68%DF/ F, p<0.0001) **F.** (Top) Day 1 Mean conditioning trial-by-trial responses (bottom) zoom on CS-evoked responses and US-evoked responses. **G.** CS-evoked and **H.** US-evoked trial-by-trial Area Under Curve computed among for mice. **I.** Day 2 Mean Ca2+ signal retrieval time course. **J.** Mean Ca2+ signal during BL vs. CS vs. expected-US stimulation. **K.** Day 2 Mean retrieval trial-by-trial responses **L.** CS- and US-evoked trial-by-trial AUC during Retrieval

On Habituation Day VTA^VGluT2+^ neurons were insensitive to the CS as shown in the example of mouse #141 (**Fig. 2.B-C, left**), demonstrating that a neutral stimulus was insufficient to evoke a significant response in this neuronal population. During the Conditioning Day, the response of individual mice to CS was larger compared to Habituation Day. All mice had a strong evoked response to the US, similar to the unconditioned footshock experiment (**Fig. 2.B-C, middle**). Following conditioning, increased sensitivity to CS was evident in the mean responses of all mice (**Fig. 2.D; n=8**), where CS-evoked and US-evoked response amplitudes were significantly higher than the baseline (BL-evoked=0.12%DF/F; CS-evoked=3.8% DF/F, p<0.0001; US-evoked=8.68% DF/F, p<0.0001; **Fig. 2.E**), and US-evoked signal was still significantly stronger than CS-evoked signal (p<0.01). As a control, another group of mice infected with a GFP virus followed the same aversive conditioning protocol, and did not exhibit Ca^2+^ signal variations at the time of either CS and US (**Sup. Fig. 2.B-C**), demonstrating that the signal is not due to artifacts.

Furthermore, the amplitude of the CS-evoked peak of Ca^2+^ response increased gradually over trials (**Fig. 2.C, middle**). Analysis of the mean peak CS-evoked Ca^2+^ signal on a trial-by-trial basis shows an approximately linear increase across trials (**Fig. 2.F bottom: ‘CS Time’**), whilst the equivalent mean US-evoked response remained constant across trials (**Fig. 2.F bottom: ‘US Time’**). The AUC shows that CS-evoked responses (**Fig. 2.G**) increase trial-by-trial, reaching statistical significance using a t-test when comparing the first and last trial together (Trial 1=-11.2 AUC vs. Trial 5=98.9 AUC, P<0.05), while US-evoked signals remained constant (**Fig. 2.H**). GFP signal in control mice did not show any significant variation across trials (**Sup. Fig. 2.D-E**). Lastly, during Retrieval Day, the CS and expected-US signal appeared to evoke a response in VTA^VGluT2+^ neurons, as shown in the individual mouse example (**Fig. 2. B-C, right**). Mean calcium signal from all mice (n=8) reveals a CS-evoked response increase, with a rebound of activity at the time of expected-US (**Fig. 2.I**). Both CS-evoked and Expected-US-evoked response amplitudes were significantly higher than the baseline (BL=0.18 vs. CS=5.26, t-test p<0.0001; BL vs. US=2.86, p<0.005; **Fig. 2.J**), indicating that VTA^VGluT2+^ neuronal responses to conditioned aversive stimulation were sustained over time. This also strongly suggests that responses may be due not only to the presence of physical stimuli but may also be due to a component of expectation. Across successive trials, there was a trend for both CS-evoked and Expected-US-evoked peak responses to decrease by a small fraction (**Fig. 2.K & Sup. Fig. 2.A**), although this was not significant (**Fig. 2.L**). Testing for a signal decrease over trials would be interesting to investigate in the future, but the results here indicate that aversive memory is robust and persistent for at least a period of 24 h.

In summary, these results show that during Conditioning Day, the amplitude of the CS-evoked responses of VTA^VGluT2+^ neurons gradually increased over repetitions, while US-evoked activity remains relatively constant. During Retrieval Day CS-evoked responses are similar to those observed during Conditioning Day and there is an Expected-US-evoked response, which indicates that the VTA^VGluT2+^ population is sensitive and adapts to aversive conditioning and responds to aversive stimulation for at least 24 h.

### 3. VTA VGluT2 neurons respond to successive repetition of unconditioned rewarding stimulation

Since direct stimulation of VTA^VGluT2+^ neurons promote reward (Wang et al. 2015), and that unitary electrophysiological recording has shown that some VTA^VGluT2+^ neurons are sensitive to rewarding stimulation (Root, Estrin, and Morales 2018), we next wanted to investigate glutamatergic neuron population responses to rewarding stimulation. To answer this question, calcium signals were recorded whilst mice were head-fixed and a tube directly delivered a liquid reward in their mouth (5 % sucrose water, ITI=60-120 s, 30 trials) (**Fig. 3.A**). Fiber position was carefully verified in brain slices for each individual mouse at the end of experiments (**Fig. 3.B**). Following rewarding stimulation, a rapid increase of calcium activity was observed in VGluT2 neuronal populations of individual mice, as shown in **Fig. 3.C**. An increase of calcium activity just after reward delivery was observed across all mice that was significantly different from baseline measurements for 8.16 s (**Fig. 3.D**). Analysis of the amplitude of the peak of activity revealed that it was significantly higher than the baseline (BL=0.07 % DF/F vs. Reward=4.72 % DF/F, p<0.0001). To check for a potential gradual change of signal amplitude across stimulation, we calculated the mean in groups of 5 consecutive trials and found stable activity across trials (**Fig. 3.F-G**), similar to responses to unconditioned aversive stimulation.

**Figure 3.**
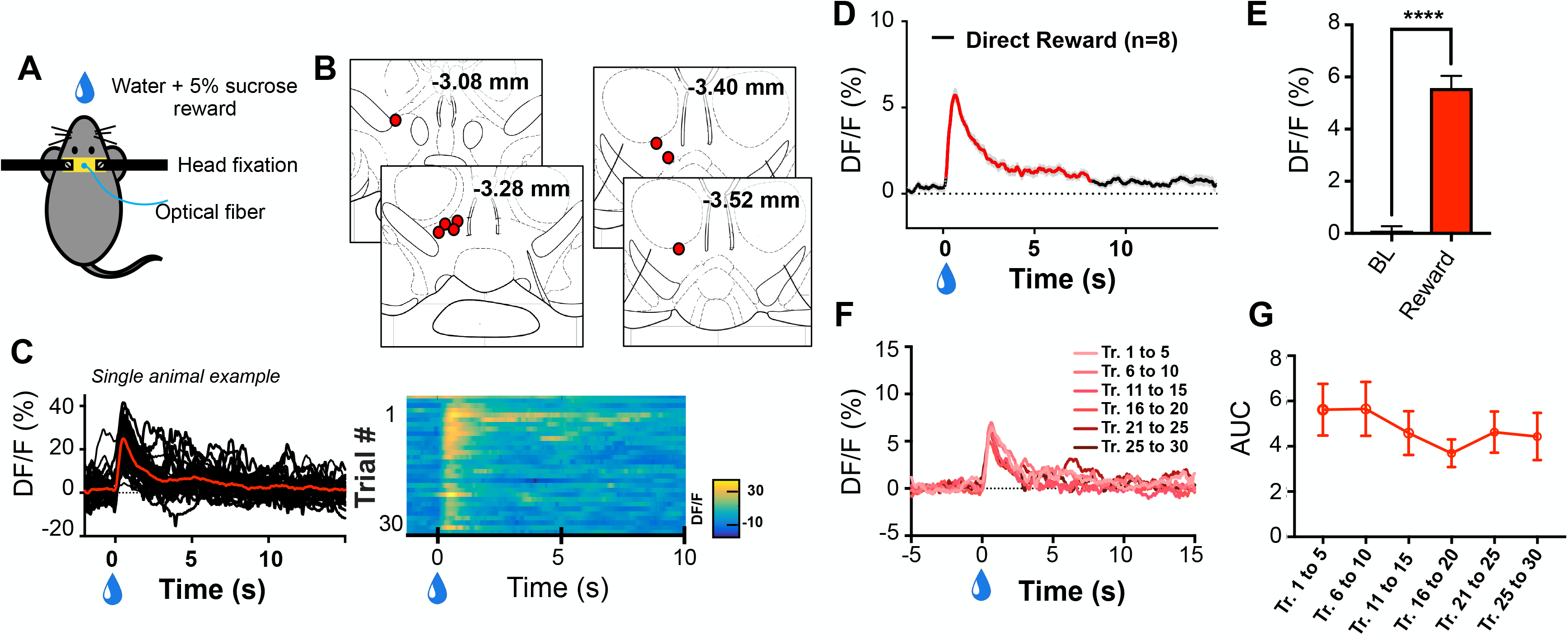
VTA response to unconditioned reward. **A.** Schematic representation of Fiber Photometry setup during reward delivery. Head-fixed mice directly receive 5% sucrose in their mouth. **B.** Location of optical fiber tip for all animals. **C.** Single animal example Ca2+ signal (left) plotted as individual time courses in black (30 trials), mean Ca2+ signal in red; (right) and plotted as heatmap for 30 trials. Water drop represents rewarding stimulation time (T=0). **D.** Mean Ca2+ signal time course (n=10 mice), increasing during rewarding stimulation. **E.** Averaged Ca2+ signal during the baseline (BL=0.07% DF/F) vs. rewarding stimulation (Reward=4.72 %DF/F, p<0.0001). **F.** Mean reward-evoked 5-trial block Ca2+ signal responses. **G.** Mean reward-evoked AUC of 5-trial blocks.

Here, we show that VTA^VGluT2+^ neurons were excited by rewarding stimulation and the amplitude of their activation remained stable across stimulation trials.

### 4. VTA VGluT2 neurons respond to reward conditioning

We have demonstrated that the VTA^VGluT2+^ population gradually learned to respond to a CS tone preceding aversive stimulation. To determine if rewarding stimulation can evoke a similar response pattern, mice were conditioned to a reward; the CS was a 5 s tone and the US was 5% sucrose water (random ITI 60-120 s). The experiment was conducted over three consecutive days, similarly to aversive stimulation described above: Habituation Day, Conditioning Day, and Retrieval Day (**Fig. 4.A**). During the Conditioning Day, an increase in calcium signal evoked by CS was observed, followed by a higher amplitude increase evoked by the US (**Fig. 4.B**). Group data confirmed this pattern (**Fig. 4.D**), revealing that the difference between baseline and CS stimulation-evoked VTA^VGluT2+^ population response was statistically significant and sustained for 3.88 s, and that the US stimulation evoked a larger activity increase that slowly returned to baseline level over 13.88 s. The CS-evoked and US-evoked activity peaks were significantly higher than the baseline (BL=0.02 % DF/ F, CS=2.53 % DF/F, p<0.0001; US=11.88 % DF/F, p<0.0001, **Fig. 4.E**), and the US-evoked activity significantly higher than the CS activity (p<0.0001). To investigate trial-by-trial changes, we plotted groups of 5 trials (**Fig. 4.F**); this shows that CS-evoked activity slowly increased across trials, which was confirmed by calculating the area under curve (**Fig. 4.G**), whereas US-evoked responses remained stable across trials (**Fig. 4.F and Fig. 4.G**). In addition, it appeared that across conditioning, CS-evoked activity remained sustained until the beginning of the rewarding stimulation, which can be seen by increasing the time window to take into account the long sustained activity during CS (Trial 1-5= 0.84 AUC vs. Trial 26-30=3.81, p<0.05, **Sup. Fig. 4.B**). During the Retrieval day, a calcium signal peak was visible during CS and US as shown in **Fig. 4.C**, but of low amplitude compared to baseline. Group mean calcium signal changes show that the CS did not evoke any clear change, whereas expected-US evoked very brief activity trend (**Fig. 4.H**); however, this was not statistically significant when looking at a 0.5 s time window around activity peak (**Fig. 4.I**). In line with these results, no pattern was found across trials (**Fig. 4.J and Sup. Fig. 4.C**).

**Figure 4.**
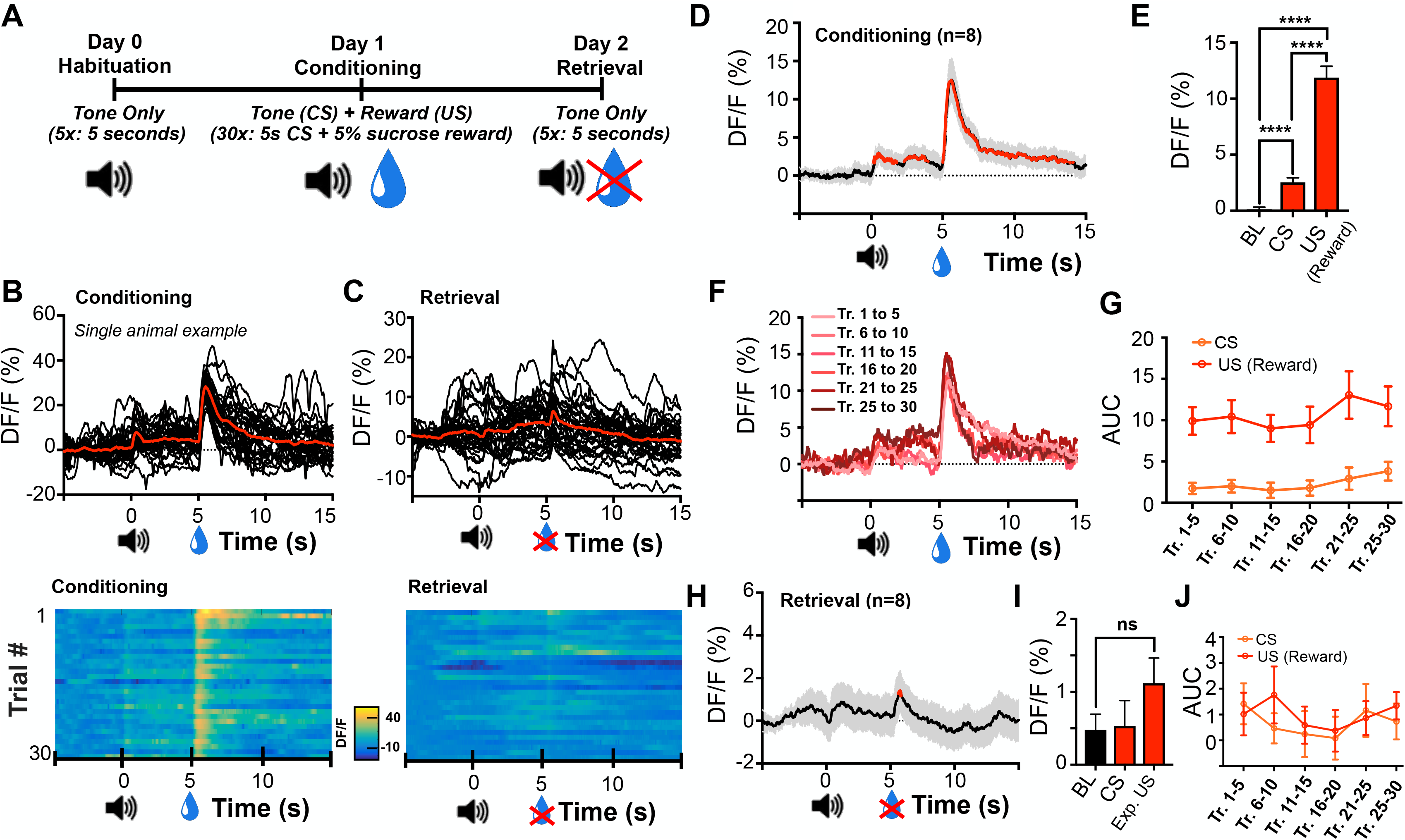
VTA response to rewarding conditioning. **A.** Schematic representation of CS-US conditioning, where CS is a 5 s tone and US a 5% sucrose liquid reward; Symbols represent tone CS and US reward time (respectively T=0s and T=5s). **B.** Single mouse example signal plotted as individual time courses in black (30 trials), and mean in red, (top) for Day 1 Conditioning (CS-US), (bottom) and plotted as heatmap. **C.** Same as panel B, for Day 2 Retrieval (CS only), (bottom) and plotted as heatmap. **D.** Day 1 Mean conditioning time course (n=10 mice), increasing during CS and US stimulation. **E.** Mean Ca2++ signal during baseline (BL=0.02%DF/F) vs. (CS=2.53%DF/F, p<0.0001) vs. (US=11.88%DF/F, p<0.0001; CS vs US p<0.0001). **F.** Day 1 Mean 5-trial block Ca2+ signal conditioning responses. **G.** Mean CS-evoked and US-Reward-evoked trial-by-trial AUC of 5-trial blocks. **H.** Day 2 Mean Retrieval time course. **I.** Mean 5-trial block Ca2+ signal during BL (BL=0.47%DF/F) vs. CS (CS=0.53%DF/F) vs. Expected Reward stimulation (Exp. US=1.12%DF/F). **J.** CS- and Expected Reward-evoked AUC.

Together, these data demonstrate that the VTA^VGluT2+^ population responds to rewarding conditioning with increasing CS-evoked activity amplitude and duration, whilst US-evoked activity remained stable. But during retrieval, VTA^VGluT2+^ neurons did not respond. These results indicate VTA^VGluT2+^ population is sensitive to rewarding conditioning, but we could not demonstrate formation of rewarding memories.

### 5. VTA VGluT2 neurons may be characterized by their specific network

The VTA^VGluT2+^ neuronal population can respond to both aversive and rewarding stimulation and is considered a heterogeneous population. Determining which broader network they belong to may be an alternative strategy to characterize them (Morales and Margolis, 2017). First, to determine whether VTA^VGluT2+^ neurons send collaterals to structures serving a similar function, we injected in the same Vglut2-cre mice both the retrograde tracer Cholera Toxin-B conjugated with Alexa 594 (CTB-594) in NAc and Cholera Toxin-B conjugated with Alexa 488 (CTB-488) in LHb, two structures downstream of VTA^VGluT2+^ neurons that are associated with aversive responses. Mice VTA were then were infected with AAV-DIO-GFP virus (**Fig. 5.A**). After expression, CTB was retrogradely transported from NAc and LHb terminals to VTA cell bodies (**Fig. 5.B-C**), confirming previous studies in the literature (Taylor et al., 2014; Faget et al., 2016). We found 28.99 CTB-594-positive neurons in VTA (Anterior=10.33; Middle=9.66; Posterior=9; **Fig. 5.D**), and 2.66 CTB-488-positive neurons (Anterior=2.66; Middle=2; Posterior=0.66), revealing that NAc and VTA receive VTA projections. The distribution of CTB along anteroposterior axis of VTA revealed both NAc and LHb received homogeneous projection from VTA^VGluT2+^ neurons (**Fig. 5.D-E**). Only a minority of labeled neurons coexpressed VGLuT2 and CTB marker (0.11% for NAc, and 0.04% for LHb), revealing glutamatergic neurons account for a minority of VTA projections to NAc and LHb. Importantly, there was no cell in VTA expressing both CTB-494 and CTB-488 (**Fig. 5.B**), suggesting that VTA^VGluT2+^ neurons do not send collaterals to these two aversive-response associated downstream VTA targets. But nine cells expressing simultaneously CTB-594 and CTB-488 were found, demonstrating that VTA non-glutamatergic neurons can send collaterals to NAc and LHb.

**Figure 5.**
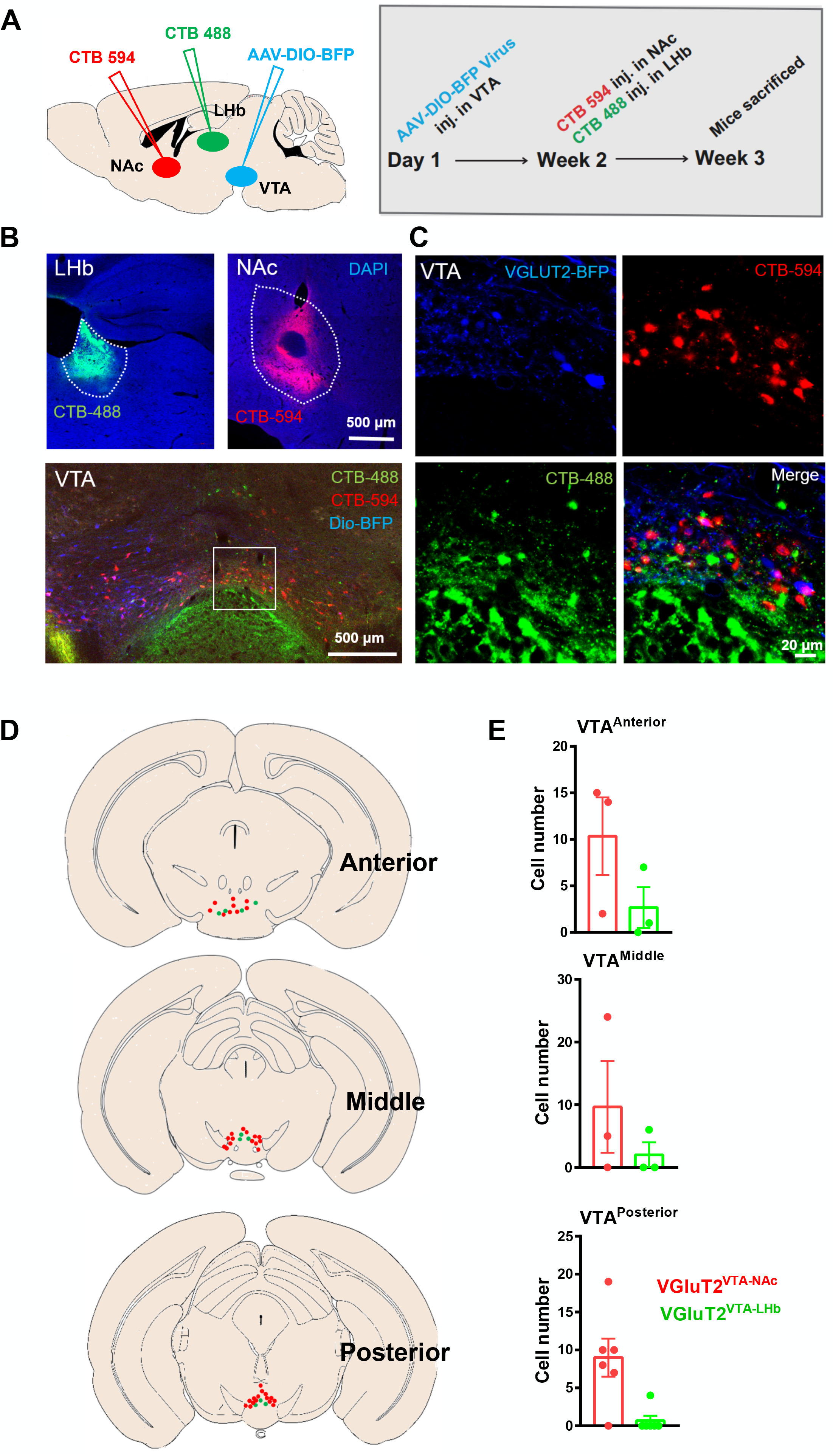
LHb and NAc projections to VTA VGluT2 neurons. **A.** Schematic representation of NAc CTB Alexa 594 tracer (CTB 594) injection (in red) + LHb CTB Alexa 488 tracer (CTB 488) injection (in green) + AAV9-DIO-BFP injection into VTA of VGluT2-cre animals. **B.** Representative images showing the injection site of CTB 488 (top left, in green) in LHb, CTB 594 (top right, in red) in NAc. AAV-DIO-BFP virus expression in VTA (bottom, in blue) and retrogradely labelling CTB cells in VTA (scale bar, 200 μm). **C.** Magnified pictures of VTA neurons expressing each fluorescence, merged on the bottom right. No co-expression of CTB 488 and CTB 594. **D.** Representative pictures of VGluT2 positive neurons (in blue) VGluT2-VTA-NAc neurons (in red) and VGluT2-VTA-LHb neurons (in green) along anteroposterior VTA axis. **E.** Quantification of VGluT2-VTA-NAc and VGluT2-VTA-LHb cells along anteroposterior VTA axis.

Finally, to check and detail the VTA^VGluT2+^ network, we mapped the structures projecting to VTA^VGluT2+^ neurons using Cre-dependent monosynaptic retrograde tracing. VGluT2-ires-Cre transgenic mice received AAV-CAG-DIO-histo-TVA-GFP (AAV2/9) and AAV-CAG-DIO-RG (AAV2/9) virus injection into VTA. Three weeks later, VTA was infected with RV-EvnA-DsRed (EnvA-pseudotyped, G-deleted and DsRed-expressing rabies virus) using the same coordinates (**Sup. Fig. 5.A**). Mice were sacrificed one week after this second injection and injection sites were verified as VTA (**Sup. Fig. 5.B**). Neurons projecting to VTA^VGluT2+^ neurons were defined as expressing red retrogradely label virus only (**Sup Fig. 5.D**), allowing a precise count of cells in each upstream target (**Sup Fig. 5.C**). VTA^VGluT2+^ upstream projections were particularly strong from the Dorsal Raphe Nucleus (DRN, 293.3%), Lateral Hypothalamus (LH, 166.4%), and the Medial Habenula (MHb, 166.5%). The Lateral Habenula (LHb, 129.7%), Rostromedial Tegmental Nucleus (RMTg, 107.2%), Laterodorsal Tegmental Nucleus (LDTg, 101.9%), and Periaqueductal Gray (PAG, 181.93%, including vlPAG, lPAG and dPAG) were also particularly strong, as well as NAc (122.6%, including NACshell and NACcore).

Together, these data show that, on one side, a majority of projections to VTA^VGluT2^ arise from DRN, LH and MHb. On the other side, VTA^VGluT2+^ neurons project to NAc and LHb, which represents only a minority of VTA projections to these structures. While VTA send collaterals to NAc and LHb, they do not originate from VTA^VGluT2+^ neurons. This may indicate that VTA-NAc and VTA-LHb VGluT2 neurons belong to two segregated networks.

## Summary of Results

We used fiber photometry to investigate how the Ventral Tegmental Area glutamatergic neuronal population was recruited by unconditioned and conditioned aversive and rewarding stimulation. We demonstrated that VTA^VGluT2+^ population was activated by both aversive and rewarding unconditioned stimulation, with a response amplitude remaining stable across trials. During the conditioning protocol, CS-evoked responses gradually increased over trials, briefly for aversive conditioning and in a sustained manner for rewarding conditioning; in parallel, US-evoked activity remained stable. During a retrieval test, CS-evoked and expected-US neural activities remained strong only for the aversive conditioning protocol, but not for the rewarding protocol. This suggests that aversive and rewarding conditioning signals are integrated by VTA^VGluT2+^ neurons through different mechanisms. Finally, to help better characterize VTA^VGluT2+^ neurons based on their connectivity pattern, we injected a CTB retrograde tracer in LHb and NAc nuclei, which revealed that only VTA non-glutamatergic neurons send collaterals to NAc and LHb. In parallel, by injecting rabies retrograde tracer in VGluT2-cre animals, we identified that VTA^VGluT2+^ neurons received inputs from variety of brain structures, with especially strong inputs from DRN, LH and MHb.

## Discussion

### Fiber photometry and neuronal response

We used fiber photometry to perform a systematic exploration of VTA^VGluT2+^ neurons at the population level. This method allowed us to selectively record the activity of VTA glutamatergic neurons by injecting a cre-dependent GCaMP6s virus in VGluT2-cre mice. Photometry is a recording method used to index synchronous neuronal activity at the proximity of the fiber tip, indirectly made through measures of intracellular variations of calcium concentration (Resendez and Stuber, 2014). Interpretation of calcium signal is not trivial since different intracellular variations can account for a fraction of the recorded signal; for example, even sub-threshold voltage events can generate calcium local increase (Engbers 2014; Perez-Reyes, 2003). Although the GCaMP6s signal mainly corresponds to soma response (Chen et al., 2013), it has been demonstrated that changes at the level of the terminal fields can also be detected (Gunaydin et al., 2014). Consequently, we cannot rule out that part of the signal we recorded may originate from terminals of locally infected VGluT2 VTA neurons, forming synapses with neighboring dopamine or GABA neurons at the microcircuit level. But such local interactions may represent only a small proportion of the signal detected and their contributions are negligible. Finally, unlike single neuronal recordings, a strong variation in calcium signal may be not only indicative of the activity of a population but can also be interpreted as a function of number of active neurons. An increase of fluorescence may mean that more neurons are recruited by a neuronal process.

But VTA^VGluT2+^ being a heterogenous population, use of both single neuron and population recording have disadvantages. Looking ahead, it will become very useful to combine photometry with technologies such as electrophysiology (Kim, 2017) to allow simultaneous investigation of single unit and population responses. This strategy could become a key method used to understand neural network function.

### VTA glutamatergic neurons response to aversive and rewarding conditioning

This study focused on the population response of VTA^VGluT2+^ neurons during unconditioned and conditioned aversive or rewarding stimulation. We have demonstrated that: (I) VTA^VGluT2+^ exhibits a similar response to unconditioned aversive and rewarding stimulation and (II) for conditioned aversive and rewarding stimulation, during Training Day, CS-evoked activity gradually increased over trials, and US-evoked activity remained constant. The key finding here is the difference in terms of conditioned evoked responses during the Retrieval Day. For aversive conditioning, the amplitude of the CS-evoked responses increased gradually over trials, and there was a significant response at the expected time of footshock delivery. This behavior was not observed for reward conditioning.

By demonstrating that the VTA^VGluT2+^ population responds to both to rewarding and aversive stimulation, our results are in line with optogenetic studies showing VTA^VGluT2+^ neurons can promote both rewarding behaviors when directly stimulating VTA (Wang et al., 2015), and aversive behaviors when stimulating VTA^VGluT2+^ downstream targets (Qi et al., 2016; Root et al., 2014). These opposing functions may be explained if, on one hand, VTA^VGluT2+^ projecting neurons encode aversion, whilst on the other hand, VTA^VGluT2+^ local neurons encode reward. Another hypothesis is that VTA^VGluT2+^ neurons encode perceived saliency, like recently demonstrated in the paraventricular thalamus (Zhu et al., 2018). But further investigation is required to account for the scalability of response with stimulus intensity.

Our data also complements another recent electrophysiological study showing that VTA^VGluT2+^ neurons are sensitive to both aversive and rewarding stimulation (Root, Estrin, and Morales, 2018). This provided a precise characterization of individual VGluT2 neurons based on their individual response pattern: they revealed that most VGluT2 neurons increased firing rate during aversive stimulation, and decreased during rewarding stimulation (Root, Estrin, and Morales, 2018). In addition, they also showed that the majority of VTA^VGluT2+^ neurons decreased their activity during reward, and only a small fraction was activated by both reward and aversion. Our results are in accordance with and support their findings that VTA^VGluT2+^ neurons increase activity during unconditioned aversive stimulation. Another of their findings, that most of VTA^VGluT2+^ neurons decreased activity during reward, appears at odds with our results. However, this can perhaps be explained by the fact that the GCaMP6s signal is mainly correlated with neuronal activity, and it is virtually insensitive to inhibition (Chen et al., 2013). Consequently, the increase of calcium signal we recorded during reward was likely driven mainly by the small VTA^VGluT2+^ subpopulation described as responsive to both aversion and reward (Root, Estrin, and Morales, 2018), and thus may not reflect the other subpopulation that is inhibited by reward. However, further investigation combining population and single neuron recording is required to test this hypothesis.

This study next asked, for the first time, what the VTA^VGluT2+^ neuronal response to aversive conditioning and its retrieval are. We demonstrated that CS-evoked population response gradually increased across trials, whilst in parallel, the US-evoked response remained stable. We also demonstrated that VTA^VGluT2+^ neurons strongly respond to CS and expected-US 24 h after aversive conditioning. Some of these VTA^VGluT2+^ features, in particular, the gradual increased response to CS over trials, and response to an expected stimulation, may resemble VTA dopamine neurons (Nakahara et al., 2004; Satoh et al., 2003; Bromberg-Martin et al., 2010; Okihide, 2009; Nasser et al. 2017). One possible explanation our results arises from the finding that a subpopulation of VTA glutamtergic neurons, VGluT2 neurons, can coexpress VGluT2 and TH (Yamaguchi et al., 2011; Root et al., 2014; Morales and Margolis, 2017). We cannot exclude that our results for the CS-evoked response is due to VGluT2 neurons expressing TH. Another non-exclusive explanation may be that DA and VGluT2 neurons form connections at the microcircuit level (Dobi et al., 2010), which could modulate the response of VGluT2 neurons during CS and US pairing, especially modulating the VGluT2 population response during CS.

Finally, CS preceding-reward or CS preceding-aversive responses are slightly different, the former being sustained and the later brief, suggesting that each of these CS-evoked responses is integrated by a different subpopulation across trials. Supporting this idea, and contrary to the aversive experiment result (Fig. 2.I), the retrieval-evoked response to the CS, following rewarding conditioning, remained extremely weak, if not absent (Fig. 4.H). This suggest that VGluT2 neurons responding to reward and aversive conditioning may belong to segregated subpopulations, probably in part corresponding to the different types of VTA^VGluT2+^ neurons recently characterized (Root, Estrin, and Morales, 2018). These divergences may be in part explained by specific connectivities of VTA^VGluT2+^ neurons sensitive to reward and aversion, in particular at the microcircuit level where interactions with dopamine and GABA neurons are known to exist (Dobi et al., 2010). For example, we can hypothesize than local VTA^VGluT2+^ neurons encode rewarding functions, while VTA^VGluT2+^ projections encode aversive ones. Future experiment should compare the functional and response profiles of these VTA^VGluT2+^ neurons that have diverging connectivity patterns.

Together, we showed that across trials, more and more VTA^VGluT2+^ neurons are similarly recruited during rewarding and aversive unconditioned stimulation, but differences emerge during conditioning. In particular, retrieval responses diverged, suggesting that neurons responding to reward and aversion conditioning may belong to different subpopulations. Investigating the response of VTA^VGluT2+^ neurons during conditioning could help understand VTA function, including mechanisms that sustains learning and expectation in DA and GABA neurons.

### Understanding VTA^VGluT2+^ neurons based on their network

We used CTB retrograde tracing to investigate VTA^VGluT2+^ projections to NAc and LH and found that VTA^VGluT2+^ represent a minority of projections to these structures. Of particular importance, we showed that, whilst VTA sends collaterals to NAc and LHb, these collaterals do not arise from VTA glutamatergic populations. Knowing that both VTA^VGluT2+^-to-NAc and VTA^VGluT2+^-to-LHb pathways are known to serve aversive function (Root et al., 2014; Wang et al., 2015), our results raise the question of their individual functional characteristics. In particular, it would be important to separately record VTA^VGluT2+^-to-NAc population response and VTA^VGluT2^+-to-LHb, for example, by using fiber photometry at the terminal level, to compare their activity patterns during aversive stimulation.

We next used RV tracing to map the inputs of VTA^VGluT2+^ neurons and observed that particularly strong projections to VTA glutamatergic neurons were coming from DRN, LH and MHb. Our data are consistent with previous studies (Taylor et al., 2014; Faget et al., 2016), and confirm that structures such as LHb or NAc also sends projections specifically to VTA^VGluT2+^ populations, which may supply feedback that promotes aversive behaviors. It is known that DRN and PAG send projections to VTA-DA and VTA-GABA neurons linked to aversive and rewarding behaviors (Qi et al., 2014; Ntamati, Creed, and Luscher, 2017; Ntamati et al., 2018); however, we observed that these structures also send parallel projections to VTA^VGluT2+^, whose function remains unknown.

The diversity of VTA^VGluT2+^ neurons inputs and outputs support the idea that VTA^VGluT2+^ function is not only based on their molecular background, but also on the network the belong to (Morales and Margolis, 2017). For example, functions and response patterns of dopamine neurons are highly heterogeneous and could depend of their specific projecting pattern (Morales and Margolis, 2017; Lammel et al., 2011; Lammel, Lim, and Malenka, 2014). In particular, knowing VTA^VGluT2+^ neurons stimulation can be either aversive (Root et al., 2014; Wang et al., 2015) or rewarding (Wang et al., 2015), we can posit that VTA^VGluT2+^ projections promote aversion, while local VTA^VGluT2+^ neurons promote reward, likely via neighboring connections to DA and GABA neurons. An interesting future direction would be to specifically target these potential subpopulations based on their connection patterns at the circuit or microcircuit level, to investigate and systematically compare their activity profile and their molecular background.

In summary, we used fiber photometry to demonstrate that VTA^VGluT2+^ neuronal population response to aversive and rewarding conditioning are divergent, especially during retrieval of conditioning. This suggests that VTA^VGluT2+^ populations responding to reward or aversive conditioning may belong to different subpopulations. In the future, investigating VTA^VGluT2+^ neurons based on their local or long-range connectivity pattern may be important to better understand VTA function. In particular, deciphering the function of VTA^VGluT2+^ at the microcircuit level would shed new light on our understanding of how local VTA DA and GABA neurons process reward and aversive conditioning.

## Acknowledgments

We thank Ji Hu for providing the VGluT2-Cre mice. This work was sponsored by the National Natural Science Foundation of China under grant numbers NSFC 31630031 (L.W.), 81425010 (L.W.), NSFC 31671116 (J.T.) 31500861 (Z.Z.) and 31471109 (L.L.); I n t e r n a t i o n a l P a r t n e r s h i p P r o g r a m o f C h i n e s e A c a d e m y o f S c i e n c e s 172644KYS820170004 (L.W.); External Cooperation Program of the Chinese Academy of Sciences GJHZ1508 (L.W.); Guangdong Provincial Key Laboratory of Brain Connectome and Behavior 2017B030301017 (L.W.); CAS President’s International Fellowship 2017PB0090 (M.Q.); Shenzhen governmental grants JCYJ20150529143500959 (L. W.), JCYJ 20160429190927063 (J. T.), Shenzhen governmental grants KQJSCX20160301144002 (Z.Z), JCYJ20150401150223647 (Z.Z.), and JSGG20160429190521240 (F.Y.); Shenzhen Discipline Construction Project for Neurobiology DRCSM [2016]1379 (L.W.) and Ten Thousand Talent Program (L.W.).

## Contributions

M.Q., Z.Z., Z.L. and X.L. contributed equally to this work. M.Q. and Z.Z. designed and initiated the project. Z.Z. performed virus injections and fiber implantation. Z.L., P.Z., and K.H. setup the behavior protocol. Z.L., P.Z., and C.C performed photometry experiments. Y.L., C.C. and K.H. helped to collect the data. M.Q. and SL.P. processed and analyzed photometry data. X.L., Z.L. P.Z. and Y.L performed immunohistochemistry and quantitative analyzed of the tracing data. M.Q. and Z.Z. interpreted the results. M.Q, ZZ, X.L., SL.P. and L.W commented the manuscript. M.Q., Z.Z. and L.W wrote the manuscript. L.W. supervised all aspects of the project.

---

**Supplementary Figure 2.**
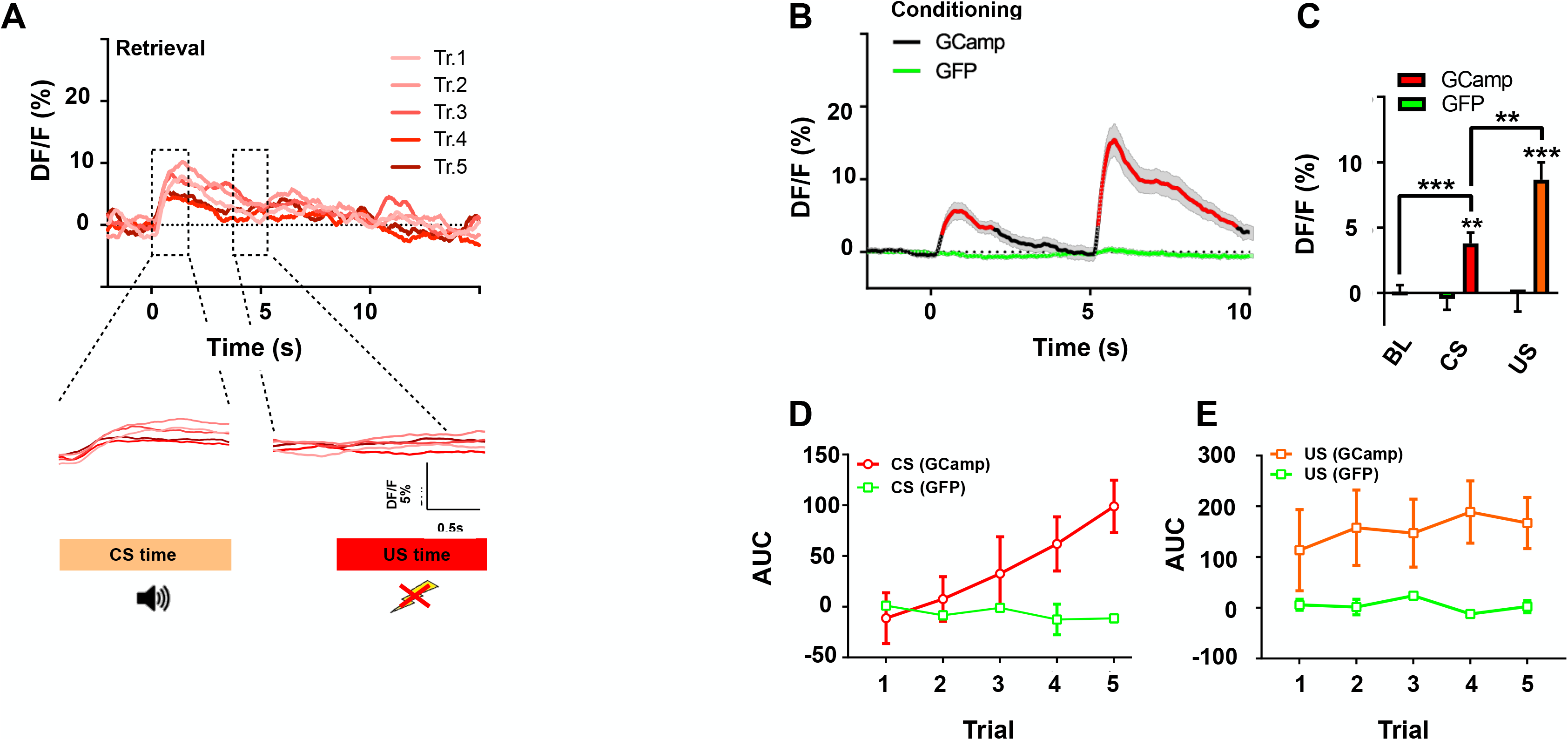
VTA response to aversive conditioning. **A.** Day 2 Mean retrieval trial-by-trial responses including (bottom) magnified CS-evoked responses and US-evoked responses. **B.** Day 1 Conditioning time course including control GFP animals (n=3 mice), remaining flat. **C.** Mean Ca2+ signal same as figure 2.E, during BL vs. CS vs. US, including GFP animals. **D.** Trial-by-trial AUC for CS-evoked response combining figure GCaMP mice from Fig. 2G in red, and GFP animals in green **E.** Trial-by-trial AUC for US-evoked response same as Fig. 2.H, including GFP mice in green.

**Supplementary Figure 4.**
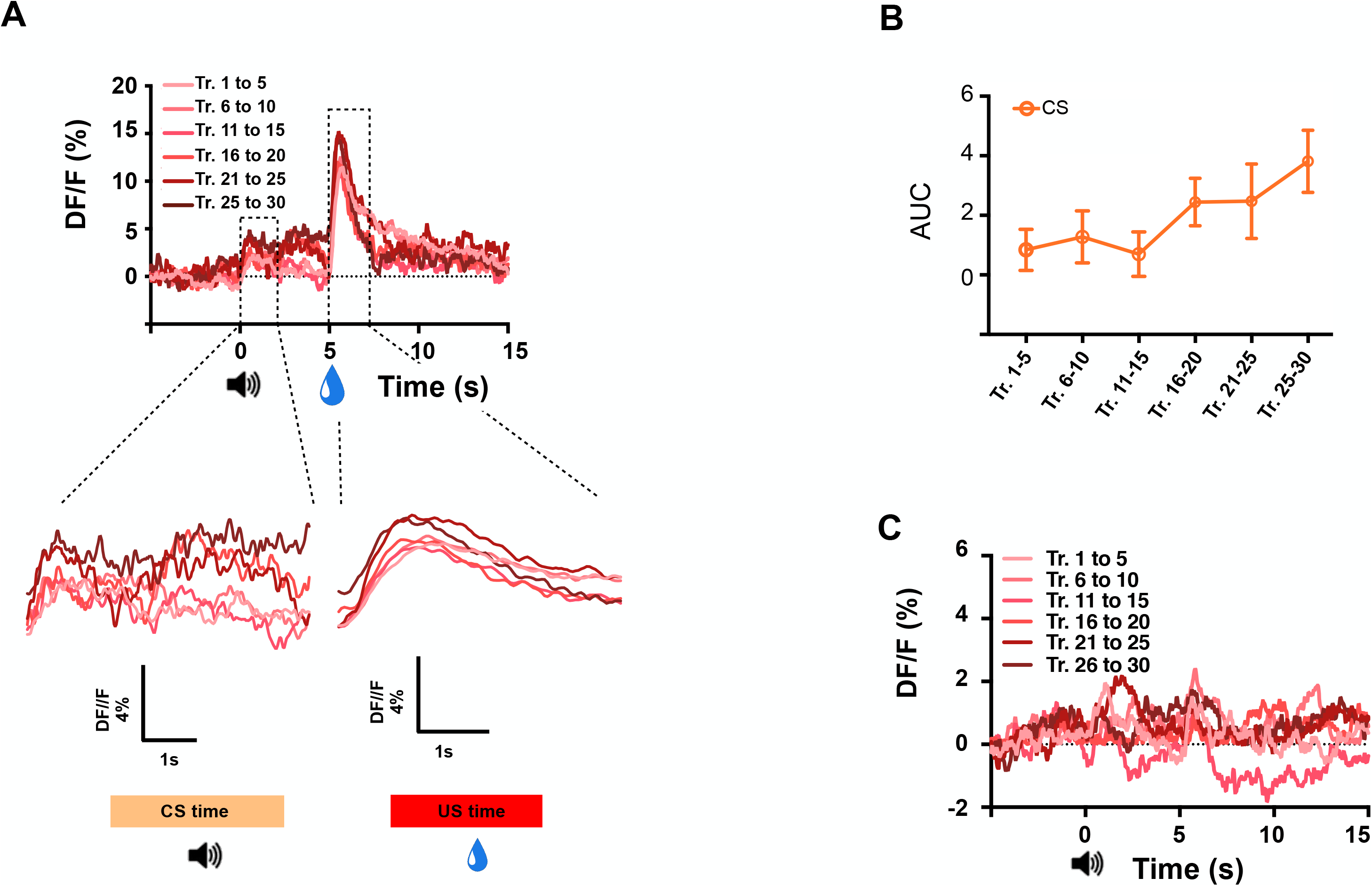
VTA response to rewarding conditioning. **A.** Day 1 Conditioning trial by trial responses averaged among all animals including (bottom) zoom on CS-evoked responses and US-evoked responses. **B.** CS-evoked trial by trial AUC, averaged by groups of 5 trials, among all animals, using a time windows including the sustained calcium signal ([0s:5s]) **C.** Day Retrieval (CS only) responses averaged by groups of 5 trials, among all animals.

**Supplementary Figure 5.**
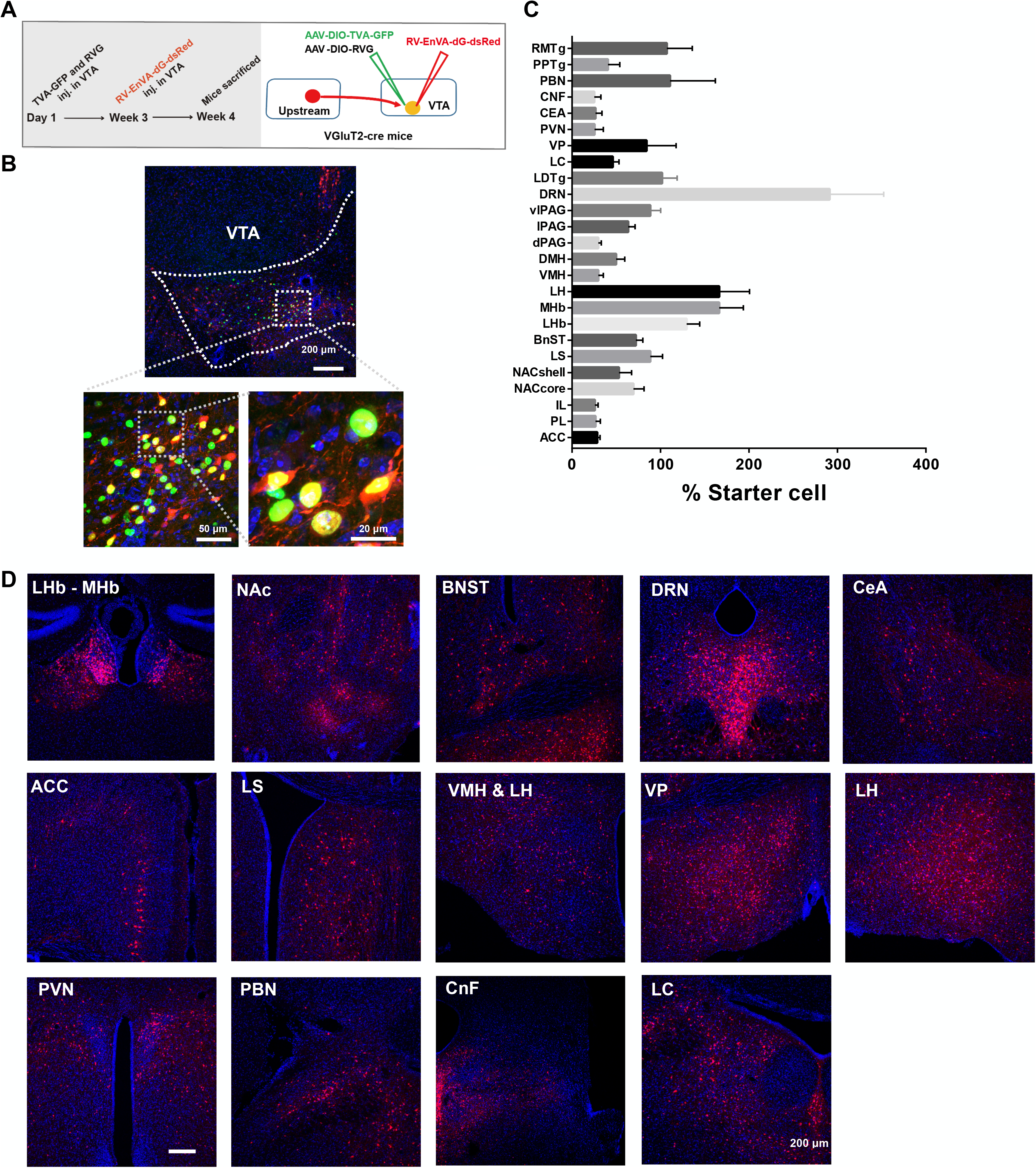
VTA VGluT2 neuronal inputs. **A**. Schematic representation of RV-virus injection protocol. **B.** Representative picture of VTA neurons expressing RV virus after injection in VTA of VGluT2-cre animals **(Red, rabies-dsRed; green, TVA-GFP; blue, DAPI, scale bar, 200 μm, 50 μm and 20 μm representively)**. Boundaries of VTA are drawn in white dashed lines. **C.** Quantification of neurons expressing RV virus after injection in VTA of VGluT2 animals. **D.** Representative pictures showing retrograde labeling in the LHb & MHb; BNST and NAc with inputs to VTA VGluT2+ neurons (Red, rabies-dsRed; blue, DAPI, scale bar, 200 μm). **ACC**, Anterior Cingulate Cortex; **BnST**, Bed nucleus of the Stria Terminalis; **CEA**, Central nucleus of the Amygdala; **CNF**, Cuneiform Nucleus; **DMH**, Dorsomedial Nucleus of the Hypothalamus; **DRN**, Dorsal Raphe Nucleus; **IL**, Infralimbic Cortex; **LC**, Locus Coeruleus; **LDTg**, Laterodorsal Tegmental Nucleus; **LH**, Lateral Hypothalamus; **LHb**, Lateral Habenula; **LS**, Lateral Septum; **MHb**, Medial Habenula; **NAC**, Nucleus Accumbens; **dPAG**, Periaqueductal Gray; **lPAG**, lateral Periaqueductal Gray; **vlPAG**, ventrolateral Periaqueductal Gray; **PBN**, Parabrachial nucleus; **PL**, Prelimbic Cortex; **PPTg**, Pedunculopontine tegmental nucleus; **PVN**, Paraventricular Nucleus; **RMTg**, Rostromedial Tegmental Nucleus; **VMH**, Ventromedial Hypothalamus; **VP**, Ventral Pallidum.

